# Relationships of diversity and evenness in adaptation strategies of the effect of protective coloration of animals

**DOI:** 10.1101/2021.05.06.441914

**Authors:** Yu. Bespalov, P. Kabalyants, S. Zuev

## Abstract

This study aims to investigate the relationship between the diversity and uniformity of the protective coloration of animals. The protective coloration of animals is the result of the formation of adaptation strategies. The aim of this study is to identify the colorimetric parameters that characterize the adaptive properties of the protective coloration of animals in relation to their natural habitat. The relevance of this study is related to the importance of the tasks of remote collection of information about various biological species; in particular, animals that carry dangerous infections. This is all the more important in light of the current epidemic problems facing humanity. The revealed colorimetric characteristics can be used to describe digital images and their further recognition. They can be classifying features in machine learning algorithms. We give an example of digital image processing of a duck (*Anas platyrhynchos)*. The analysis of the distribution of the found colorimetric parameter makes it possible to identify the image segments corresponding to the *Anas platyrhynchos*. The results thus obtained are also of interest from the point of view of the problem of the adaptive role of the diversity of biological systems in relation to the problem of the mechanisms of functioning of the protective coloration of animals.

## Introduction

The coloration of animals is a very important, topical subject of research in the field of fundamental biology and ecology [1-3], as well as in a range of high-level interdisciplinary research [4-7]. It must be said that in modern conditions the results of these fundamental studies can be of great practical importance. More specifically, this means the following. In the period of global climatic changes, the disturbance of the ecological balance in many cases will produce threats to biosafety. The spread of dangerous infectious diseases is a good example of these threats. This includes epidemics and pandemics that create extreme situations. In this regard, the practical importance of ф rapid gathering of the information on the dynamics of the number and migration of animals–potential and real natural reservoirs and carriers of the causative agents of these diseases–is increasing. Along with these animals, other representatives of the fauna, e. g., their natural enemies and competitors, become important as a subject of research. In extreme situations of epidemics and pandemics, information on resources of commercial organisms playing a certain role in ensuring food security will be demanded. (In such countries as the USA, Canada, Russia, this information, in particular, concerns the objects of pasture fish farming in regions with a low population density.)

The role of remote (aerospace) methods of collecting such information on vast and often hard-to-reach areas is increasing. The protective coloration of animals (PCA) is an obstacle to the effective use of such methods. Accordingly, the methods for image processing for eliminating the masking effect of PCA are needed. These methods can be developed with the use of results of the study of adaptive strategies of the effect of PCA. This work aims at studying the aspects of these adaptation strategies resulting from the diversity and evenness of PCA.

### The diversity of colorimetric parameters as an aspect of adaptive strategies of the effect of protective coloration of animals

The adaptive strategies (ASs) of the effect of PCA that have arisen during biological evolution provide a possibility to camouflage in an animals’ habitat. These ASs are subject to certain patterns. A formalized description of these patterns makes it possible to develop methods for automatic remote identification of the presence of animals in certain areas of terrains. In such a context, this could involve other approaches to solving the problem of eliminating the masking effect of PCA, e. g., approaches focused on image processing that facilitates such identification by visual methods.

The diversity of PCA plays an important role in the implementation of these ASs. This closely tied to the importance of this diversity for disruption of the integral visual perception of a silhouette, when the animal is situated between an observer and the background on which the silhouette is visible. In such a case, as stated in [1, 2], edge sections of the silhouette play a great role. The diversity of PCA in these areas should provide a sufficiently high probability of color fusion of some fragments of PCA with the background.

It is about the background of actual habitats, the colorimetric parameters (CPs) of which have a certain variability in time and space. The diversity of PCA provides the animal’s adaptation to this variability.

There may be cases when ASs of the effect of PCA is been implementing due to the destruction of the integral visual perception of characteristic details of the animal’s color pattern. The details (spots, stripes, etc.) on any part of the body are meant. This is possible due to the color fusion of the mentioned details of the pattern with objects located between the animal and an observer (bushes, uneven soil, etc.). In both cases, the use of the term “camouflage disguise” for ASs of the effect of PCA would seem correct in accordance with the existing precedents [1, 2]. Further, for the sake of simplification, we will mainly talk about the color fusion of fragments of PCA with the background, in cases where it is permissible in the context to neglect the differences between such fusion with the background and fusion with objects on the ground between an observer and the animal. In similar contexts, it is about CPs of the background.

Within the area of fundamental biology, the problem of the adaptive significance of the diversity of PCA can be considered as a part of a more general problem. Specifically, a more general problem of the influence of the diversity of biological system on the character of its performance (in particular, on the stability of some important aspects of the effect, up to the protection of the system from destruction, e. g., eating of an individual by a predator). Regarding ecological systems, approaches based on the Shannon-Weaver index [8-12] have received significant development in the study of this problem. For further consideration, it should be noted that these approaches involve the use of not only the diversity itself–the number of objects that compose the system (communication channels, biological species, trophic chains, etc.). Along with the diversity expressed by such a measure, the degree of evenness of values of certain measures of manifestation of performance of these objects (passing through the communication channels of amounts of information, biomasses of species, energy in trophic chains, etc.) matters.

The diversity of PCA should correspond to the diversity of CPs of habitats. However, for a variety of reasons, in a large number of cases, the diversity of CPs of habitats will be greater than that of PCA. In many cases, it is about the diversity of CPs of vegetation cover. These CPs have certain dynamics (e. g., seasonal), which often differs significantly from the dynamics of CPs observed for PCA. (The latter dynamics may be practically absent.) In this regard, within the framework of this paper, the problem of a formalized description of the indicated dynamics arises. For the case of PCA, this may be about a formalized description of non-dynamic aspects of the performance of the system, adapted to dynamics of CPs of the vegetation cover.

In mentioned extreme situations of epidemics and pandemics, they will often be faced with a lack of time and resources, in particular, technical means for collecting environmental information. Accordingly, it will be important to describe the dynamics of the state of biological objects on the basis of actual data that have a number of drawbacks. Specifically, relatively small data arrays with gaps that do not reflect the specified dynamics in real time in sufficient detail may be accessible. (E.g., data can be obtained from digital cameras on light drones, in conditions of obstacles for regular filming due to bad weather, poor visibility, etc.) Below, the results that illustrate possibilities of a formalized description of ASs of the effect of PCA based on the precedents of using the data with such lacks are described. It is about the results presented in [13, 14, 15] on a formalized description of the CPs’ dynamics for relatively simple plant communities. These results were obtained by using advanced mathematical models [16, 17]–Discrete Models of Dynamical Systems (DMDS). It will also be about modeling non-dynamic aspects of the ASs of the effect of PCA with the use of DMDS. Specifically, it is about aspects that allow an animal to adapt to the dynamics of plant communities of habitats. More specifically–to all aspects of general diversity associated with this dynamics, both in time and in space, of CPs of the vegetation cover of habitats. Among others, the aim of the current study is these aspects associated with relationships of the diversity of colorimetric parameters of PCA with their evenness.

### Study of adaptation strategies with the use of diversity of colorimetric parameters of protective coloration of animals involving DMDS

The DMDS model provides a formalized description of the structure of relationships between components of the system under study. The structure is calculated on the basis of correlations between these components. (So, initial data may not reflect directly the dynamics of the system’s states in real time.) The DMDS model provides a formalized description of the structure of inter-component and intra-component relationships caused by positive and negative influences of components on each other [17]. On the base of this structure, for certain initial conditions, an idealized trajectory of the system (ITS) can be calculated. The trajectory represents a cycle of change of system’s states; the values of components are expressed in conditional scores [18].

In [18], these combinations were called “*strategies-combinations*” (SCs); their number was considered as a measure of the system’s diversity. Specifically, the arsenal of strategies that the system has to adapt to the variety of conditions of its performance was considered as a measure of diversity.

The method of spatio-temporal modeling called “re-synchronization” is proposed [18]. It makes it possible to construct an ITS reflecting the dynamics of the system in time on the basis of the reading of the state of the system’s different parts at one moment of time. Re-chronization exploits the assumption that different parts of the system change their state by one cycle, but at a moment of time, they have different phases of the trajectory. These phases correspond to different conventional time steps, in which different SCs are observed. Different combinations of values of the system’s components correspond to different SCs.

A method similar to re-synchronization enables, on the basis of data on brightness combinations in different parts of the spectrum on different parts of the animal’s body, to calculate an idealized pseudo-trajectory of the system (IPTS). This IPTS is specific for an AS of the effect of PCA. As a rule, this is about the ratio of values of the brightness of different parts of the spectrum to the total brightness of all these spectral parts. An IPTS can be considered as an ITS of the dynamics of changes in CPs for a certain (real or hypothetical) plant background, to the color diversity of which the ASs of the effect of ASs would be optimally (conditionally, ideally) adapted in space and time. For calculating both the ITS and IPTS, the correlation matrix between the system components are used. In this case, the values of the brightness of different parts of the spectrum at different parts of the animal’s body are components. The ratios of these brightness values in these different places are interpreted as SCs that provide camouflage pattern in terrain areas having a certain set of CPs. The dynamics of SCs in real time do not matter in connection with such a use of a correlation matrix. (For the case of PCA, the dynamics are often missing.) Therefore, the IPTS can be considered as a visual representation of a set of SCs that provide the effect of PCA. That is about providing performance due to a rather high probability of color fusion with the background of the fragments of PCA with one or another SCs determined by the ratio of the brightness in different parts of the spectrum. That is about the fusion color with objects on the ground (bushes, uneven soil, etc.) located between the animal and an observer, but not with the background. In the first case, the biological meaning of ASs of the effect of PCA is the destruction of the integral visual perception of the silhouette of the animal against the background of a certain area of habitat. In the second case, this meaning is the destruction of the integral visual perception of characteristic details (spots, stripes, etc.) of the picture of PCA.

As a measure of diversity can be adopted a number of different SCs in the cycle for both IPTS and ITS. Regarding the subject of our consideration, this means the diversity of the set of strategies of the effect of PCA. That is about a diversity of color combinations of PCA’s fragments. Specifically, that is about the diversity providing a certain probability of color fusion of some fragments of PCA with the background or objects on the ground located between the animal and an observer. As mentioned above, this fusion provides the effect of destroying the integral visual perception of the silhouette or the characteristic details of the PCA pattern. Notice that such a measure of diversity may not necessarily be calculated with the use of values of all system components taken into account in the modeling involving DMDS.

In [15, 18, 19, 20, 21], abilities to use the differences between the background’s ITS and the animals’ IPTS in the methods of image processing for improving the detection conditions on the ground are shown. In [22], such abilities based on the results of comparisons of relationships’ graphs between color parameters of PCA and the plant background constructed by DMDS are shown. In particular, this was shown for visual detection on processed digital pictures. From the point of view of practical use of results presented in [15, 18, 19, 20, 21], it is important that initial data for them was obtained by image processing of RGB-pictures of animals and their habitats. These pictures can be taken from the board of widespread models of light drones by equipment that is included in the standard delivery set. In the aforementioned extreme situations, the available fleet and such aircrafts can be mobilized for collecting actual data for informational support of decision-making. In [13, 14, 23], the ability to use similar results for the development of methods for remote assessment of other biological factors that are important from the point of view of emergence and elimination of biosafety threats. This is about the factors not considered directly in this work. Specifically, it is about the factors associated with the mass reproduction of toxic cyanobacteria (there is a good example in the Baltic sea) and possible technologies for preventing eutrophication, which creates this threat to biosafety.

SCs can have enough complicated combinations of brightness values in different ranges of the spectrum. Comparison of the ITS of the plant background and the IPTS built for PCA allows one to find its SCs that have some diversity. This set of SCs, to some extent, corresponds to ASs of the effect of PCA corresponding to the widest possible range of variability of background CPs in time and space. At the same time, there are factors that determine the difference in the degree of color diversity of PCA and the background, in many cases, with a wide diversity of the background. Analysis of differences in the diversity of background CPs and SCs of the effect of PCA creates certain prerequisites for new approaches to the development of image processing methods. This is about methods eliminating the masking effect of PCA with the use of system colorimetric parameters. Specifically, SCPs reflecting certain aspects of diversity, in particular, numerical values of its certain measure, are considered. Differences between these aspects for CPs of the surrounding area and PCA enable to eliminate the masking effect of PCA. The results of applying such an approach are presented in [18, 20].

In [19], they considered the case when the general color diversity of habitats cannot be described using DMDS and presented in the form of ITS. It is about the diversity of CPs of urban habitats of the gray rat (*Ratus norvegicus Berk*). In this case, one can proceed from the premise that the diversity of PCA is likely less than the diversity of CPs habitats in whole. But at the same time, the diversity of PCA will be higher than the diversity of CPs of individual parts of habitats with high probability. Therefore, it makes sense to seek an attribute of PCA that provides the maximum diversity of the set of SCs. According to results of [19] obtained with the help of DMDS, the expression R/(R + G + B)*G/(R + G + B) can correspond to such an attribute of PCA. The results of digital processing of the picture of a rat against the background of a fragment of the urban landscape presented in [19] point that the use of this attribute is promising for the elimination of the masking effect of PCA of this biological species. This species is able to create serious threats to biosafety. The results in **[20]** are about the usage of similar expression R*G/(R+G+B) for eliminating the masking effect of PCA of a gray rat and toad aga (*Rhinella marina*). In this case, it is about a masking effect against the background of plant communities in habitats of these biological species. The diversity in time and space of CPs of these plant communities can be modeled with the use of DMDS and resynchronization and presented in the form of ITS. Similar results on the base of the PCA data of a representative of ichthyofauna–pike *(Esox lucius)* are presented in [18]. The product of (R + G)/(R + G + B) by R/G was used.

In a certain way, the above facts show the role of diversity in the ASs of the effect of PCA. To understand the specific conditions of such an implementation, it makes sense appropriate to involve some provisions of the concept of optimal diversity presented in [12]. This concept was developed primarily for ecological systems and highlights the importance of the availability of certain resources for the creation of optimal diversity.

For PCA, the angular size of a silhouette in certain observation situations (e. g., when a predator is hunting for an animal, or an animal is hunting for its prey) can be such a resource. The amount of this resource restricts the number of multi-colored spots (significantly different in their CPs) having a sufficiently large angular size on the silhouette. For this size, there will be no visual fusion of adjacent spots. Biochemical or genetic factors [4-7] predicted by A. Turing model can also act as such a limiting resource [4]. It is important for further consideration that there is a limitation on the diversity of PCA associated with the lack of a certain resource. It is about the diversity of values CPs of fragments of PCA.

Let us consider the aspects of ASs, which make it possible to compensate for the negative effect of this limitation on the effect of PCA.

It was mentioned above that use of the Shannon-Weaver index enables to take into consideration not only the diversity of the system but also its evenness. In [3, 24], the results of the study of relationships of diversity and evenness of different animals in different conditions of their habitat on the data about PCA are presented. Notice also that these relationships can be seen in the cycle of the so-called Margalef’s model of succession [25]. Some aspects of the dynamics of this model correspond to aspects of many relatively simple plant communities. In this cycle, there are phases with different values of pigment diversity expressed by the “yellow-green index” as well as expressed by the same index, the degree of evenness of presence of green chlorophylls and yellow-orange-red plant pigments.

Further, the relationships of diversity and evenness and their role in ASs of the effect of PCA will be considered.

### Relationships of diversity and evenness and their role in adaptive strategies of the effect of protective coloration of animals

Let us consider another issue before considering the mechanisms of compensation of the influence of lack of a certain resource on the diversity PCA. Specifically, consider maximum values of parameters of this diversity corresponding to an ideal AS of camouflage. For example, for enough frequent and significant case of seeking an animal between an observer and a plant background. The variability in time and space of CPs of plant communities is determined by different ratios of the availability of green chlorophyll and other plant pigments–red, yellow, and orange–in communities. The range of the set of SCs of the effect of PCA should be suitable. Specifically, from SCs with a predominance of red tones to SCs with a predominance of green. A lack of a certain resource may hinder the implementation of a fairly wide range of such SCs. The possible causes of the appearance of such lack are mentioned above. The reason may be just the impossibility for an animal with certain physiology to produce any pigment. (Thus, mammals, in most known cases cannot independently produce green pigment. In the case of availability of the pigment, its source is other organisms, e g. sloth, in whose wool algae settle, giving it a green color, masking the animal in foliage.)

A sufficiently effective AS of the effect of PCA can be implemented by leveling the CPs. Thus, in the case of many mammals, the balanced PCA is brownish brown. Such a balanced PCA also has a certain diversity. It seems another circumstance plays an important role. Specifically, CPs of fragments of such PCA provide, not quite perfect, but satisfactory color fusion in different conditions for masking purposes. That is about conditions of detection of an animal in areas on the ground and, with some of the ground details, the color of which reveals a predominance of both green and red-yellow components. This fusion can be observed at different moments of time and points of space during the formation of CPs of plant communities. It provided the destruction of the integral visual perception of the silhouette or feature elements of the pattern of PCA for sufficient different situations.

The evenness can increase the versatility of the effect of the animal’s camouflage but at the expense of completeness of color fusion of PCA’s fragments with the background and terrain’s parts.

The question on options for the implementation of optimal combinations of diversity and evenness in ASs of the effect PCA arises.

It seems plausible to assume that the role of evenness increases with the decrease of importance value or opportunities for implementation of diversity. Here the reasons for a decrease in the diversity of PCA are considered. Notice that they quite often, but not always, are not associated with the lack of a certain resource. But they can be related, e. g., to parameters of the diversity of CPs of habitats. In such cases, in ASs of the effect of PCA, it is reasonable to supplement the decreasing diversity with an increasing uniformity. (With the reduction or prevention, actual or potential, scarcity of any resource. This supplementation can be interpreted as an aspect of the implementation of the principles of the above-mentioned concept [12] of optimal diversity.)

In [3], examples of such relationships between the roles of diversity and evenness in ASs of the effect of PCA of different representatives of Australian fauna. It is about biological species with sharp differences in the evolutionary history of their adaptation to habitats changing on different scales of time. The examples of the following species were considered: feral one-humped camel (*Camelus dromedarius*), dingo dog (*Canis lupus dingo*), and kangaroo (*<u>Macropus rufus</u>*).

Camels have become a part of Australian fauna very recently–about a hundred years ago–in terms of the rate of biological evolution. At the same time, their intensively increasing population is constantly expanding its habitat. This habitat is currently occupying new, increasing areas of Australian deserts and semi-deserts. The plant communities of these areas have enough expressed variability of their CPs in time and space. Accordingly, PCA of camels is distinguished by pronounced diversity (the largest out of three aforementioned Australian animals.) At the same time, in these three cases, the evenness of PCA of camels is the lowest. (In order to avoid confusion, it should be noted that in figures presented in [3], the ordinate is the inverse of the evenness of the red and green components of the RGB model of digital pictures of animals and the plant background of their habitats.)

The adaptation time of kangaroos to Australian habitats is immeasurably longer than that of camels and dingoes. During this adaptation, the stabilizing natural selection should bring the diversity of PCA closer to the optimal one. It is possible that this process included the decrease in diversity of PCA, compensated due to evenness. In accordance with these premises, in results presented in [3], PCA of kangaroo has the least diversity and the greatest evenness in comparison with camels and dingoes.

Dingoes have an intermediate position between kangaroos and camels in terms of their stay in Australia (thousands of years). Accordingly, as it is presented in [3], the values of diversity and evenness of PCA of dingoes have the same intermediate position in comparison with camels and kangaroos.

It can be assumed that similar relationships between the diversity and evenness of PCA in different parts of the animal’s body. In particular, it is about the areas that play different roles in the implementation of ASs of the effect of PCA, including the destruction of the silhouette’s integral visual perception. The importance in this sense of its edge sections is emphasized in the above-mentioned works [1, 2]. It can be assumed that in these areas and in others, remote from them, the roles of diversity and evenness in ASs of the effect of PCA. Accordingly, it may be assumed that the character of correlation of values of certain measures of diversity and evenness can be used for automatically identifying the presence of certain animals in a certain area of the terrain. That is about animals, the implementation of ASs of the effect of PCA of which supposes a certain character of such a correlation, moreover absent in the color of animal’s habitats.

These assumptions confirm the study of the nature of the correlations between the diversity and evenness of PCA in the framework of this work. This study was conducted on the free access digital pictures of a female mallard duck (*Anas platyrhynchos*) and habitats of this biological species. Nowadays, this species is of practical use for both hunting and its role as a carrier and natural reservoir of “bird flu” (*Grippus avium*). Fragments of these digital pictures with images of a duck’s body and the corresponding area of the habitat were divided into segments, which, in turn, were divided into microsegments. For pixels of each micro-segment, the average value of R/G (hereinafter referred to as the R/G value) was calculated. For a set of microsegments of each segment, the following statistical parameters that reflect the diversity and evenness of the R/G value were calculated:

‐ mean;
‐ range–the difference between the maximum and minimum values;
‐ standard deviation;
‐ mode;
‐ mode amplitude.

The following statistically significant (p<0.05) Spearman correlations of PCA were revealed for the duck but not revealed for the background of its habitat: positive correlations–between the mean and the range, as well as between the mean and the standard deviation; negative–between the mean and the of the mode amplitude.

This effect, regarding ASs of the effect PCA, may have the following biological meaning.

The mode amplitude reflects the size and expressiveness and, accordingly, the likelihood of appearance and identification of certain features of the pattern of the protective coloration of animals–stripes, spots, certain patterns of their placement (e. g., the frequency of stripes or spots of the most common color.) The increase of the amplitude mode can be compensated by approaching the mean to the optimum of evenness that provides a satisfactory degree of color fusion with the background (in particular, in this case, by decreasing the mean with its approaching to the optimum of evenness, located in the subset of relatively low values in the range of observed values in PCA of a duck of the R/G value.) The statistically significant negative correlation between the mode amplitude and the mean of R/G can be explained by this effect.

This approach to the optimum of evenness, due to the decrease in the mean of R/G, might also compensate for the decrease of evenness. Conversely, if diversity takes a high value, there is no need to approach the optimum of evenness by reducing the mean of R/G. Statistically significant positive correlations between the range and the mean of R/G mean, as well as the mean and standard deviation of R/G, may be explained by this.

Thus, the correlation structure of the color characteristics of PCA observed, in this case, has, apparently, a certain biological meaning and manifests some aspects of ASs of the effect of PCA. These aspects are related to relationships of coloration’s diversity and evenness.

Different statistics that characterize diversity and evenness can be interpreted as different types of resources that can be used in the ASs of the effect of PCA. In this case, the similarities to concepts proposed in [26] for describing ASs of performance of the cardiovascular system and in [27] for describing ASs of performance of a fish population can be built. That is about concepts that consider ASs with the simultaneous or sequential use of several types of resources.

The results of processing digital profile pictures of a female mallard duck in Fig. 1 show the implementation of such ASs.

**Figure 1.**
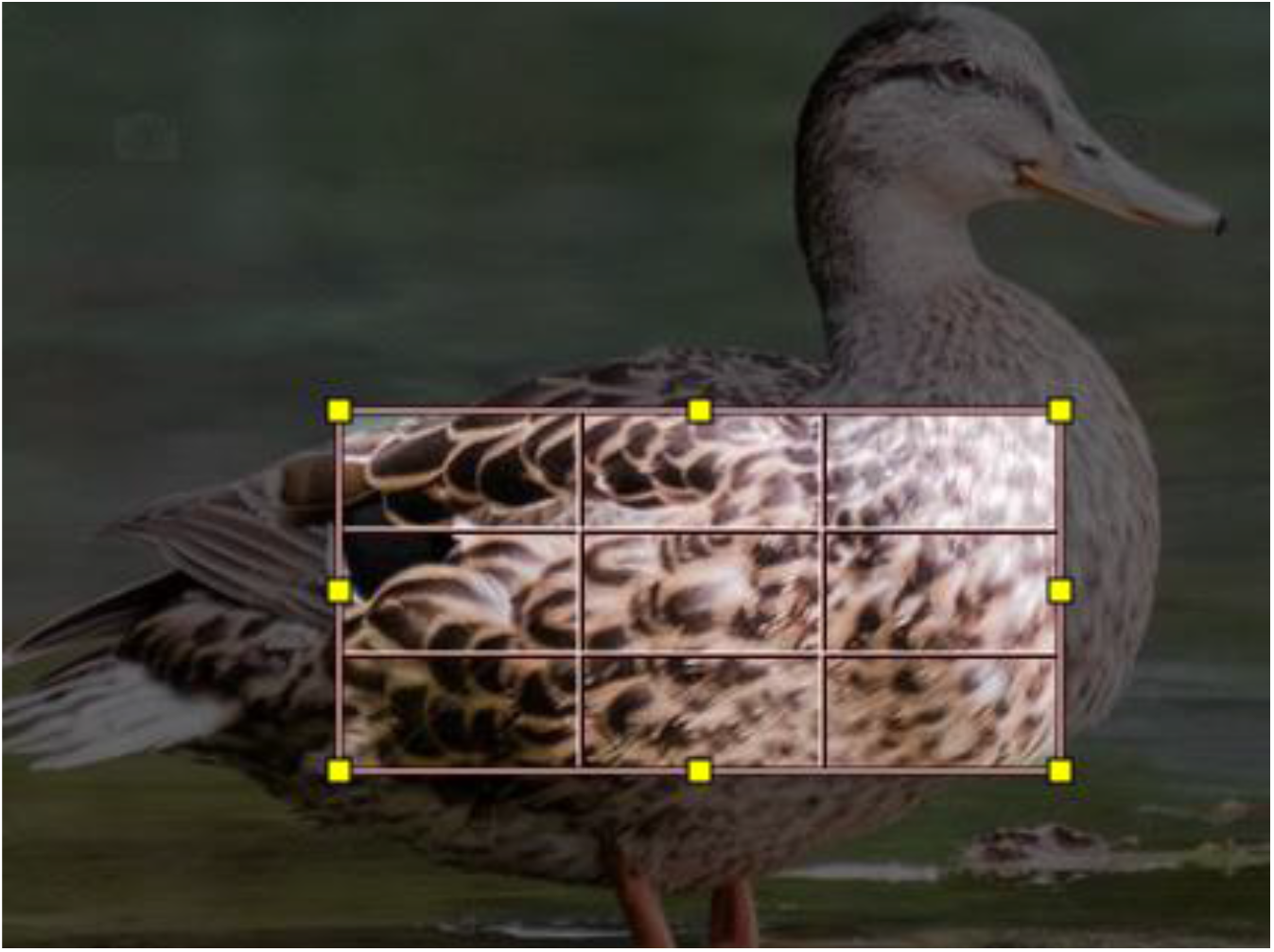
Digital picture of a female mallard duck; the rectangle corresponds to the processed part of the picture. The plots on the silhouette with different values of statistics characterizing the diversity and evenness of PCA were analyzed.

The character of the distribution of the parameters of diversity and evenness of PCA on the silhouette, presented in Tables 1-4, generally corresponds to the above correlations between these parameters. The biological meaning of such the character of distribution, apparently, can be defined as a tendency to compensate for the lack of diversity or expressed development of certain elements of its coloration that can unmask an animal due to evenness. Such a tendency can be interpreted as the implementation of ASs of the effect of PCA with alternate use of resources. At the same time, some parts of the body include a combination of low values of both diversity and evenness of PCA. That is about small parts located in such a manner that their role in ASs of the effect of PCA is apparently insignificant. More precisely, this is the case of the AS, which main biological meaning is to save several (maximum possible number) types of resources at once.

**Table 1.**
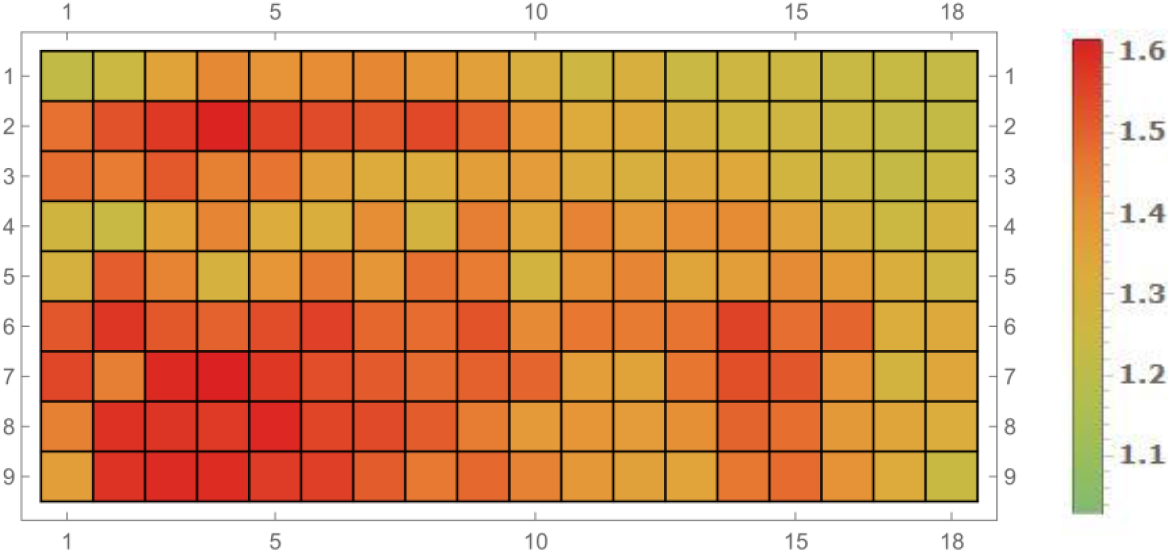
Distribution of segments of the silhouette’s part of the female mallard duck marked in Fig. 1, with different means of R/G that reflects the degree of distance from the optimum of evenness.) On the left: the conditional color scale of values. Red-brown colors on the conventional color scale correspond to large values of the mean of R/G, middle values–yellow shades, small values–shades of green.

**Table 2.**
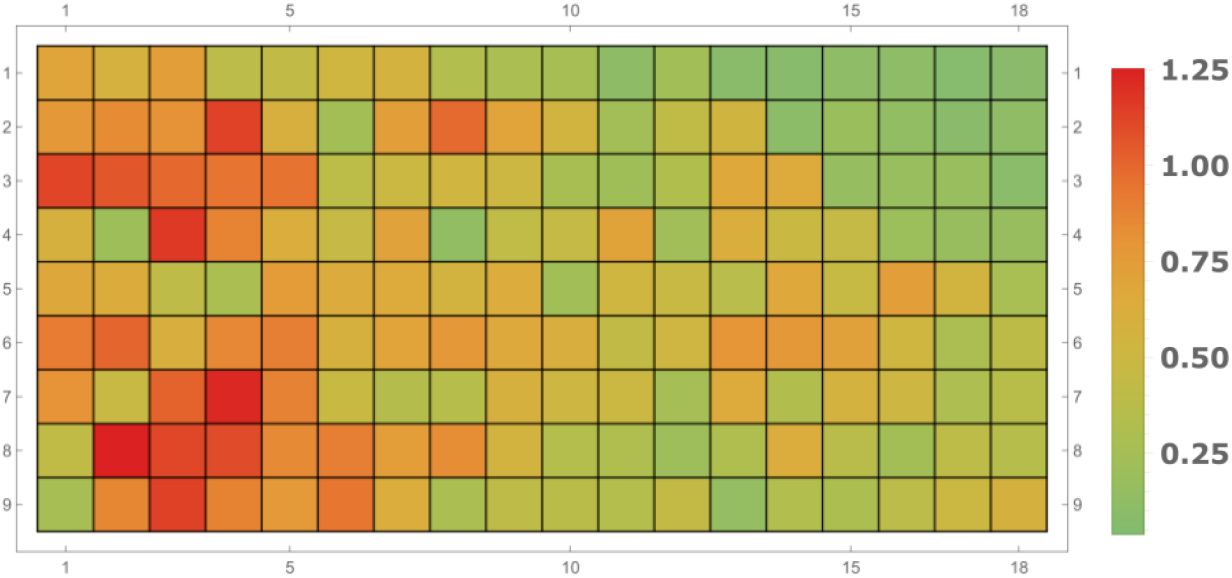
Distribution of segments of the part of the female mallard duck’s silhouette marked in Fig. 1 with different values of the range of R/G. On the left: the conditional color scale of values. On the conventional color scale, red-brown colors correspond to large values of the mean of R/G, middle values–yellow shades, small values–green shades.

**Table 3.**
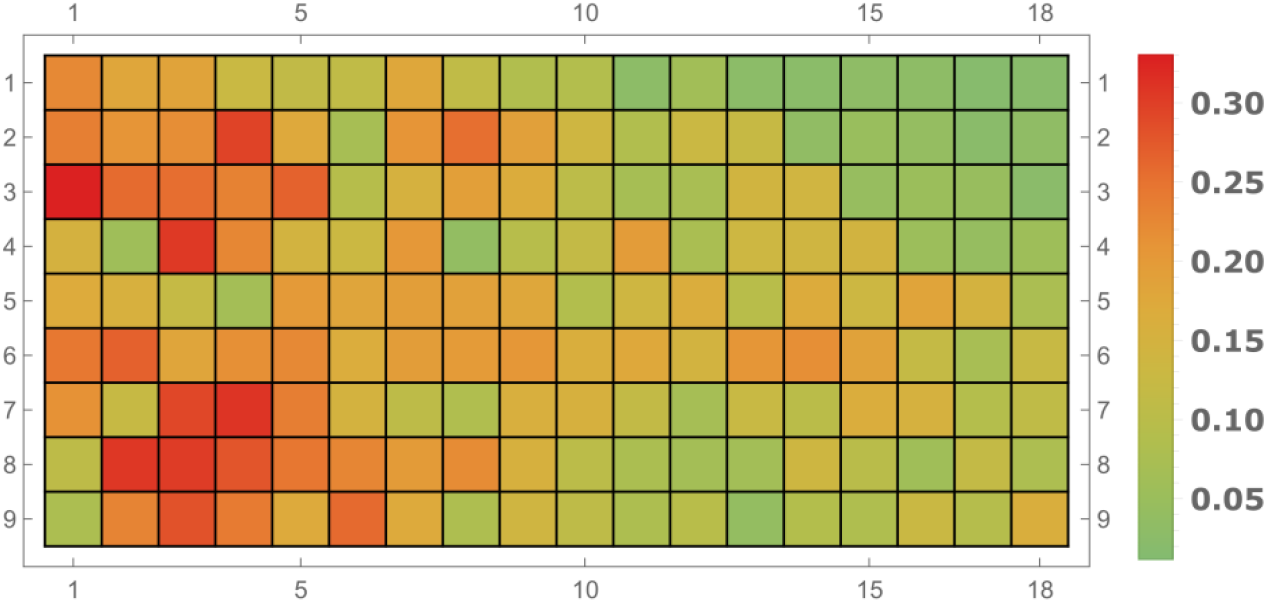
Distribution of segments with different values of the standard deviation of R/G on the part of the female mallard duck’s silhouette marked in Fig. 1. On the left: the conditional color scale of values. On the conventional color scale, red-brown colors correspond to large values of the standard deviation; of R/G, middle values– yellow shades, small values–green shades.

**Table 4.**
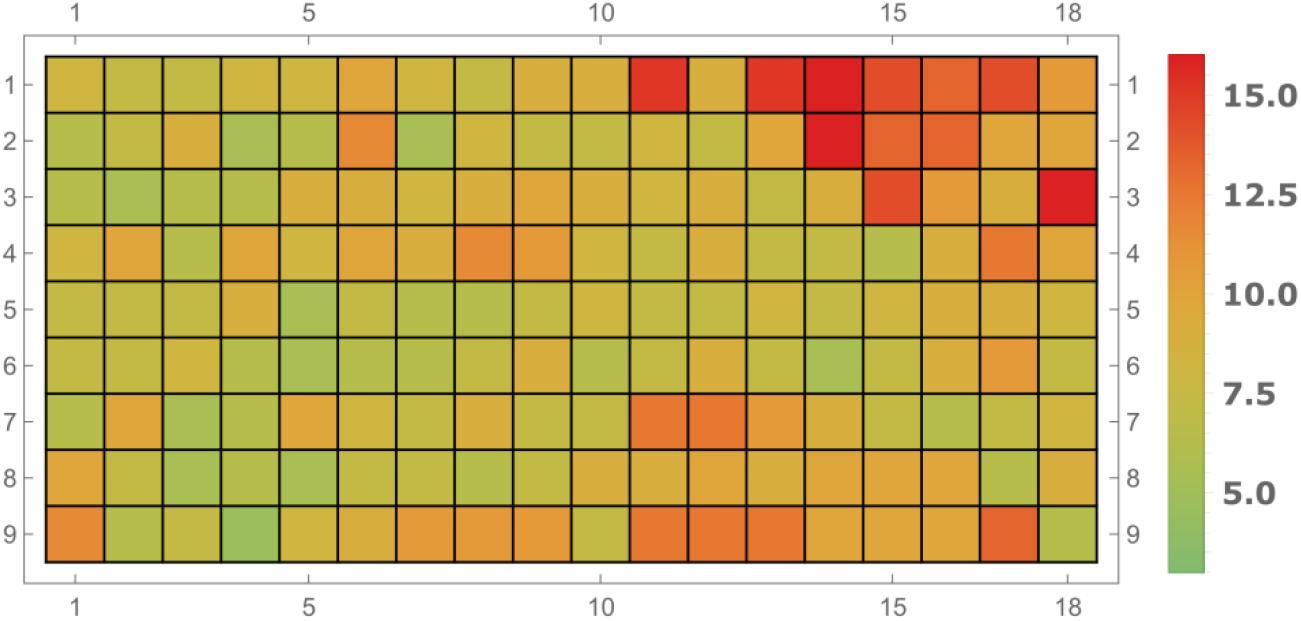
Distribution of segments with different values of the mode amplitude of R/G on the part of the female mallard duck’s silhouette marked in Fig. 1. On the left: the conditional color scale of values. On the conventional color scale, red-brown colors correspond to large values of the mode amplitude of R/G, middle values– yellow shades, small values–green shades.

The described results also have a certain practical value. It is about the possibilities of their application for the development of procedures for automatic identification of signs of the animals’ presence of a certain species on the ground. (Specifically, this is about mallard ducks; remote automatic methods of monitoring the localization and migration of this species should be demanded currently to struggle with a transfer of pathogens of dangerous infections over great distances.) These methods can also be used to eliminate interference, the sources of which are objects (bunches of grass, bumps, etc.) that resemble animals in certain colorimetric parameters, but with the absence of the above-described patterns of relationships of coloration’s diversity and evenness.

The development of such procedures with the use of neural networks seems promising.

## Discussion and conclusions

The results of the study of the role of diversity and evenness and their relationships in adaptation strategies of the effect of protective coloration of animals presented in this work, in our opinion, provide new opportunities for the development of new image processing methods having practical importance. These methods will eliminate the effects of camouflage masking the animals on the ground. In particular, these above methods can be applied for automated remote registration of the number and migration of animals–in relation to the tasks of providing biosafety. (For example, the methods may control the factors that determine the dynamics of the number of animals–real and potential carriers of dangerous infectious diseases.)

At this stage, there are some prerequisites to new approaches for the development of these methods. Specific technologies for the implementation of these methods will be developed bearing in mind the conditions of their application. The technologies with the use of neural networks seem to be promising.

On the other hand, in our opinion, obtained results are also of certain interest from the point of view of fundamental biology. It is about the study of some aspects of the problem of the role of diversity of biological systems in their performance. It is important to note that within the framework of this work, certain aspects of this problem were investigated in the context and with involving the data, which currently have a rather serious applied significance.

It also relates to some of the results, which in the framework of this work can be considered as side results. It is about the above-mentioned results [13, 14, 18] of mathematical modeling of the structure and dynamics of some plant communities (i. e., communities of photosynthetic organisms.) Mass reproduction of toxic cyanobacteria observed quite recently in the Baltic sea is a resonant example of the threat to biosafety associated with such communities. Situations, in which the elimination of this threat to biosafety will require treatment from air suppressing the biological productivity of patches of these photosynthetic organisms, are possible. These patches may be kilometers-long and moving in water areas of hundreds and thousands of square kilometers. The severity of this situation will increase repeatedly when especially toxic cyanobacterial mutants appear (naturally or as a result of the activity of bioterrorist structures.)

As such situations are possible, the methods of image processing based on the results presented in [13, 14] will be required. They will help to establish the boundaries of accumulations of toxic cyanobacteria in water areas [13] and their areas most vulnerable to certain types of air processing [14].

So, the conclusion of this work on a certain theoretical and practical significance, taking into account the above, seems to be quite grounded.

The project is funded in part by the Russian Foundation for Basic Research, under Project No. 19-29-09056.

## REFERENCES

1. Endler J. A., Mappes J. The current and future state of animal coloration research. Philosophical Transactions of the Royal Society of London B: Biological Sciences. 2017, 372, 1724.

2. Duarte, R. C. Camouflage through colour change: mechanisms, adaptive value and ecological significance / R. C. Duarte, A. V. Flores, M. Stevens // Philos Trans R Soc Lond B Biol Sci. 2017. Vol. 372(1724). doi:10.1098/rstb.2016.0342

3. Yu. Bespalov, K. Nosov, O. Levchenko, O. Grigoriev, I. Hnoievyi, P. Kabalyants. Mathematical modeling of the protective coloration of animals with usage of parameters of diversity and evenness. bioRxiv 822999; doi: https://doi.org/10.1101/822999

4. Alan Mathison Turing. The Chemical Basis of Morphogenesis // Philosophical Transactions of the Royal Society of London. Series B, Biological Sciences. V. 237. No. 641 (Aug. 14, 1952). Pp. 37–72.

5. J.D. Murray. A pre-pattern formation mechanism for animal coat marking // Journal of Theoretical Biology, Vol. 88, No. I, pages 161–199; 1981.

6. J.D. Murray. On pattern formation mechanisms for lepidopteran wing patterns and mammalian coat patterns. // Philosophical Transactions of the Royal Society of London, Series B, Vol. 295, No. 1078, pages 473–496; October 7, 1981.

7. P.K. Maini, T.E. Woolley, R.E. Baker, E.A. Gaffney, S.S. Lee, Turing’s model for biologicalpattern formation and the robustness problem. Interface Focus 2(4), 487–496 (2012).

8. Shannon, C. E. and Weaver W. (1948) A mathematical theory of communication. The Bell System Technical Journal, 27, 379–423 and 623–656.

9. MacArthur R.H. Fluctuations of animal populations, and measure of community stability // Ecology. 1955. V. 36. № 7. P. 353–356.

10. Margalef R. Information theory in ecology // Gen. Syst. 1958. №3. P. 36–71.

11. Magurran A.E. Measuring biological diversity. – Oxford, UK.: Blackwell Publishing, 2004. – 256 p.

12. Bukvareva, E. N. Optimization, Niche and Neutral Mechanisms in the Formation of Biodiversity / E. N. Bukvareva, G.M. Aleshchenko // American Journal of Life Sciences. – 2013. – V. 1. – No. 4. – p. 174-183. – doi: 10.11648/j.ajls.20130104.16.

13. Bespalov Yu., Nosov R., Kabalyants P. (2017) Modeling systemic colorimetric parameters as a tool for processing images of clumps of toxic cyanobacteria targeted at their boundaries detection. bioRxiv 232413; doi: https://doi.org/10.1101/232413.

14. Olena Vysotska, Marine Georgiyants, Kostiantyn Nosov, Yurii Balym, Anna Pecherska, Andrii Porvan, Sergey Pavlov, Victoriya Shekhovtsova, Tetiana Klochko, Andrii Solodovnikov. Development of a spatialdynamical model of the structure of clumps of toxic cyanobacteria for biosafty purposes. Eastern-European Journal of Enterprise Technologies, v. 6, n. 10 (96), p. 64–75, Dec. 2018. ISSN 1729-4061.

15. Balym Y., Vysotska O., Pecherska A., Bespalov Y. Mathematical modeling of systemic colorometric parameters unmasking wild waterfowl. Eastern-European Journal of Enterprise Technologies 2017 vol: 5 (2 (89)) pp: 12–18.

16. Bespalov Y. G., Zholtkevych, G. N., Nosov, K. V., Marchenko M. S., Marchenko G. P., Psarev V. A., Utevsky A. Yu. (2008) Study of heliobiological effects using a discrete model of dynamic systems with feedback. Proc. VIII Gamow International Astronomical Conference-School in Odessa “Astronomy and beyond: Astrophysics, Cosmology, Radioastronomy and Astrobiology”, 2008. – P. 12–13.

17. Zholtkevych, G. N., Bespalov, Y. G., Nosov, K. V., Abhishek, M. (2013) Discrete Modeling of Dynamics of Zooplankton Community at the Different Stages of an Antropogeneous Eutrophication. Acta Biotheoretica, 61(4), 449–465.

18. Bespalov Yu., Nosov K., Kabalyants P.(2017) Discrete dynamical model of mechanisms determining the relations of biodiversity and stability at different levels of organization of living matter. bioRxiv 161687; doi: https://doi.org/10.1101/161687.

19. Nosov, K. V., Bespalov, Yu. G., Vysotska, O. V., Strashnenko, H. M., Pecherska, A. I. Determination of the systemic colorimetric parameters of unmasking rats at video-registration in urban conditions. Bulletin of NTU “KhPI”. Series: New solutions in modern technologies. – Kharkiv: NTU “KhPI”, 2018, 26 (1302), 2, 22–30, doi:10.20998/2413-4295.2018.26.28.

20. Vysotska O., Balym Y., Georgiyants M., Pecherska A., Nosov K., Bespalov Y. Modeling of a procedure for unmasking the foxes during activities on the elimination of biosafety threats related to rabies. Eastern European Journal of Enterprise Technologies 2017 vol: 5 (10-89) pp: 46–54

21. Bespalov Y. G., Kabalyants, P. S., Nosov K. V. (2019) Discrete modeling of dynamic systems in the problem of remote accounting of wild animals. Science-intensive technologies and innovations: Proc. Int. Conf., dedicated to the 65th anniversary of BSTU named after V.G. Shukhova, Belgorod: BSTU, 2019. – vol. 9. -P.50–54.

22. Grigoriev A.Ya., Zholtkevych G.N., Nosov, K. V., Bespalov, Yu. G., Pecherskaya A.I. (2014) Mathematical model of systemic effects of spectral characteristics dynamics of grass cover, disclosing locust crowds. Veterinary Medicine. Inter Departmental Subject Scientific Collection. V. 98, P. 154–157.

23. Vysotskaya, E., Bespalov, Y., Betin, A. (2015) Mathematical model for remote monitoring control technology bioproduction processes in water ecosystems using floating plants. Modern European Researches, (4), 121–128.

24. Bespalov, Yu. G., Nosov, K. V., Kabalyants, P. S. (2018) A mathematical model of the effect of natural selection on adaptation forms that implemented by disruptive coloration of Taurotragus oryx. bioRxiv, doi: http://dx.doi.org/10.1101/368084.

25. Margalef R., Perspectives in Ecology Theory, University of Chicago Press, Chicago, 1968. pp.112

26. Vysotskaia E. V., Bespalov Yu. G., Rak L. I., Pecherskaia A. I., Tsapenko K. V. Modeling the dynamics of the cardiovascular system parameters coherence at different stages of the adaptation syndrome. Bulletin of NTU “KhPI”. Series: Mechanical technological systems and complexes. – Kharkiv: NTU “KhPI”, 2016, 4 (1176), 79–84.

27. Bespalov Y. Discrete Modeling Dynamical Systems That Determine the Role of Biodiversity in Different Regimes of Using Resources at Different Levels of Organization of Living Matter. BioRxiv, 2018. DOI: 10.1101/281279.

